# *De Novo* Whole Genome Assembly of the Roborovski Dwarf Hamster (*Phodopus roborovskii*) Genome, an Animal Model for Severe/Critical COVID-19

**DOI:** 10.1101/2021.10.02.462569

**Authors:** Sandro Andreotti, Janine Altmüller, Claudia Quedenau, Tatiana Borodina, Geraldine Nouailles, Luiz Gustavo Teixeira Alves, Markus Landthaler, Maximilian Bieniara, Jakob Trimpert, Emanuel Wyler

**Affiliations:** Department of Mathematics and Computer Science, Institute of Computer Science, Freie Universität Berlin, Takustr. 9, 14195 Berlin, Germany; Cologne Center for Genomics, University of Cologne, Cologne. Present address: Scientific Genomics Platforms, Max Delbrück Center for Molecular Medicine, 13125 Berlin, Germany; Scientific Genomics Platforms, Max Delbrück Center for Molecular Medicine, 13125 Berlin, Germany; Charité - Universitätsmedizin Berlin, Corporate Member of Freie Universität Berlin and Humboldt-Universität zu Berlin, Division of Pulmonary Inflammation, Berlin, Germany; Berlin Institute for Medical Systems Biology, Max Delbrück Center for Molecular Medicine in the Helmholtz Association, Hannoversche Str 28, 10115 Berlin, Germany; Institut für Biologie, Humboldt Universität zu Berlin, Philippstraße 13, 10115 Berlin, Germany; Institut für Virologie, Freie Universität Berlin, Berlin, Germany

## Abstract

The Roborovski dwarf hamster *Phodopus roborovskii* belongs to the *Phodopus* genus, one of seven within *Cricetinae* subfamily. Like other rodents such as mice, rats or ferrets, hamsters can be important animal models for a range of diseases. Whereas the Syrian hamster from the genus *Mesocricetus* is now widely used as a model for mild to moderate COVID-19, Roborovski dwarf hamster show a severe to lethal course of disease upon infection with the novel human coronavirus SARS-CoV-2.

## Introduction

The ongoing pandemic caused by the human severe acute respiratory syndrome coronavirus 2 (SARS-CoV-2) has made clear that traditional animal models such as mice and rats are not always suitable to study novel diseases and moreover, multiple animal models might be required to adequately reflect a variety of possible disease manifestations (Muñoz-Fontela et al. 2020). In fact, the importance of host factors has become strikingly evident in coronavirus disease 2019 (COVID-19), as the same virus causes disease severities that span from asymptomatic infections to severe acute respiratory distress syndrome (ARDS) and fatal multi-organ dysfunction (Guan, NEJM 2020). To solve immune mechanisms, identify putative targets of interventions and to test novel therapies and vaccination regimens animals models, ideally small animal models that reflect all presentations of COVID-19, are required (Veenhuis & Zeiss 2021; Lee & Lowen 2021). Non-transgenic mice and rats could not be productively infected with and consequently showed no weight loss or lung pathology in response to SARS-CoV-2 wildtype infection (Muñoz-Fontela et al. 2020; Bao et al. 2020; Dinnon et al. 2020; Gu et al. 2020; Hassan et al. 2020; Sun et al. 2020). In order to identify suitable models, species studied in the context of COVID-19 were chosen based on similarity in the SARS-CoV-2 receptor angiotensin converting enzyme-2 (ACE-2) predicted in silico (Devaux et al. 2021; Wu et al. 2020; Pach et al. 2020). COVID-19 research comprised non-human primates (Deng et al. 2020; Lu et al. 2020; Yu et al. 2020), cats (Shi et al. 2020), ferrets (Kim et al. 2020; Richard et al. 2020), and hamsters (Bertzbach et al. 2021) as animal models. Most small animals introduced to date show mild to moderate disease symptoms with resolving infections, including Syrian hamster (Chan et al. 2020; Imai et al. 2020; Kreye et al. 2020; Sia et al. 2020; Osterrieder et al. 2020). We previously introduced a dwarf hamster species, *Phodopus roborovskii*, as a representative model for a severe course of disease including systemic immune activation and fatal disease outcome (Trimpert et al. 2020; Zhai et al. 2021). In our study, despite very similar ACE-2 sequences amongst the three analyzed Phodopus species, Roborovski dwarf hamsters (*P. roborovskii*), Campbell’s dwarf hamsters (*P. campbelli*), and Djungarian hamsters (*P. sungorus*), only the Roborovski dwarf hamster showed a severe disease manifestation following intranasal SARS-CoV-2 infection. The natural habitat of *P. roborovskii* are the sandy deserts of Northern china, where they mainly eat seeds and insects. Within groups, amicable interactions are slightly more frequent than aggressive ones, corresponding to the overall more social interactions within Phodopus compared to e.g. Mesocricetus (golden hamster) species (Wilson et al. 2009).

The Roborovski dwarf hamster is, so far, the only non-transgenic animal that consistently develops severe disease and hyper-inflammation of the lung following infection with SARS-CoV-2 (Gantier 2021; Muñoz-Fontela et al. 2020). Clinical signs develop within the first 48 hours following infection and include drastic reduction in body temperature, substantial weight loss, forced breathing, ruffled fur and lethargy. By histopathology, massive alveolar destruction and microthrombosis are evident in the lungs of infected animals while other organs, including the brain, do not seem to be primarily involved in disease development. The rapid onset and fulminant course of pulmonary disease makes this species a valuable model to study severe courses of COVID-19 in humans and test therapies and vaccinations in the background of severe disease (Muñoz-Fontela et al. 2020; Gantier 2021).

The novelty of this model however entails a problematic lack of reagents and tools to study immune reactions and other host factors. The absence of classical tools for molecular biology makes transcriptome and proteome analyses only more important as they may help to understand molecular reasons for severe COVID-19 and could supply information that helps finding reasonable medical interventions. Prerequisite for genomics and proteomics studies is a thoroughly annotated publicly available genome. Since the closest annotated and available genome comes from a species in a different genus (*Mesocricetus auratus*, MesAur1.0), we describe here a scaffold-level genome assembly based on long and short read DNA sequencing, and annotated using RNA-sequencing from heart and lung of *P. roborovskii* animals.

## Results and Discussion

### Isolation and sequencing of genomic DNA

Genomic DNA was isolated from a whole blood sample of an animal of about seven weeks of age. From the same DNA sample, libraries were prepared for Promethion long read sequencing and Illumina short read sequencing.

### Assembly

The final assembly comprises a total of 2,078 (2,055 > 50 kb) contigs with a total length of 2.38 gb, an N50 of 25.78 mb and an L50 of 30 (Supplementary Table S1). According to *QUAST*, 99.75% of 676.47 M paired-end short reads and 99.74% of 4.13 M long reads were mapped yielding average read depths of 80 and 34 respectively. The positive effect of genome assembly polishing using the described toolchain was confirmed by the genome completeness analysis with *BUSCO*. While the raw assembly as produced by *Canu* has a completeness of 79.3%, this value was improved to 85.9% with racon, 88.8% with Medaka and finally reaches 92.0% after short read polishing with *POLCA* (Supplementary Table S2). In the final step of the analysis, the screening with *Kraken2,* three contigs remained unclassified, three were classified as bacteria (total length: 314.68 kb), 74 as human and the remaining 2028 matched either the golden hamster or mouse. Of the 74 contigs classified as human 47 passed the final BLAST check and were included in the final cleaned assembly composed of 2,078 contigs.

### Annotation

Before quality and adapter trimming and filtering, the four RNA-Seq samples had between 31.2 and 13.5 million reads of which 96.4% to 98.1% passed the preprocessing stage and between 71.6% and 87.3% were uniquely mapped to the assembly with multi-mapping rates between 9.1% and 18.3% (Supplementary Table S3). The final cleaned and curated annotation based on the prediction with the GeMoMa pipeline comprises 22,139 predicted transcripts in a total of 18,029 annotated gene loci.

## Materials and Methods

### Ethics statement on animal husbandry

Roborovski dwarf hamsters were obtained through the German pet trade and housed in IVC units (Tecniplast). Hamsters were provided *ad libidum* with food and water and supplied with various enrichment materials (Carfil). For DNA extraction and sequencing, whole blood was obtained from uninfected control animals of a SARS-CoV-2 infection trial (Trimpert et al. 2020) that was performed according to all applicable regulations and approved by the relevant state authority (Landesamt für Gesundheit und Soziales, Berlin, Approval Number 0086/20). RNA was extracted from SARS-CoV-2 infected and non-infected hamsters subject to an independent experiment (Trimpert et al. 2021) under the same permit. Briefly, anaesthetized hamsters were infected with 1×105 focus forming units of SARS-CoV-2 (variant B.1, strain SARS-CoV-2/München-1.1/2020/929) in 20 μL cell culture medium.

Animals were euthanized for sample collection on days 2 and 3 post infection as previously described (Trimpert et al. 2020). Following the 3R principle, all material used for this study was obtained from animals subject to independent animal experiments, no additional animals were used.

### Isolation of genomic DNA

100 μL previously frozen whole blood was lysed by addition of 400 μL lysis solution CBV (Analytik Jena) and 10 μL proteinase K (20 mg/ml, Analytik Jena) followed by an incubation for 10 minutes at 70 °C. Following this lysis step, another 10 μL proteinase K were added to perform an extended protein digestion for 30 minutes at 50 °C. DNA was extracted using a standard phenol/chloroform extraction with a first step of adding 1 ml liquefied TE saturated phenol (Carl Roth), gentle mixing by inverting the tube 20 times and centrifugation at 10000 g for 10 minutes. The aqueous phase was aspirated with a cut pipette tip, mixed with 1 ml phenol/chloroform/isoamyl alcohol (25:24:1, Carl Roth) and mixed and centrifuged again as stated above. Again, the aqueous supernatant was carefully removed, mixed with 1 ml chloroform (Merck) and centrifuged for phase separation. The remaining aqueous phase was mixed with 1 ml absolute ethanol (Merck) and centrifuged for 30 minutes at 15000 g for DNA precipitation.

All steps were carried out with cut pipette tips and very gentle mixing to avoid shearing of the DNA.

### RNA extraction

Lung pieces were stored in RNAlater (ThermoFisher) for about 4 hours before extraction. Afterwards, the tissue was lysed in a homogenizer (Eppendorf) in Trizol (ThermoFisher). For extraction of total RNA from whole blood, 250 μL anticoagulated (EDTA) sample was lysed by addition of 750 μL Trizol LS reagent (Thermo Fisher). RNA was purified from Trizol using the Direct-zol RNA mini kit (Zymo Research) according to the manufacturer’s instructions.

### DNA short read sequencing

For short read DNA sequencing, 1ug of DNA were sonicated (Bioruptor, Diagenod), and the Illumina TruSeq DNA nano kit applied, using a slightly modified protocol with only one cycle of PCR to complete adapter structures. Following library validation and quantification (Agilent tape station, Peqlab KAPA Library Quantification Kit and the Applied Biosystems 7900HT Sequence Detection System), sequencing was performed on an Illumina NovaSeq 6000 instrument with 2×150 paired-end sequencing.

### Long-read Sequencing on PromethION

Sequencing libraries for long-read sequencing were prepared from 2.5 μg of unsheared genomic DNA, following the protocol of OxfordNanopore’s LSK109 kit (ONT, Oxford). [https://store.nanoporetech.com/eu/media/wysiwyg/pdfs/SQK-LSK109/Genomic_DNA_by_Ligation_SQK-LSK109_-minion.pdf] DNA was end-repaired, A-tailed and purified (1x of Ampure XP beads,BeckmanCoulter). Then sequencing adapter with attached motor protein was ligated and the DNA was purified with 0.4x of Ampure beads. Quality and quantity of libraries were checked using HS gDNA 50 kb kit on fragment analyzer (Agilent) and dsDNA Qubit assay (Thermofisher). Libraries were loaded three times, 30 fmol / 30 fmol / 14 fmol per 24h, i.e. 74 fmol in total. The complete runtime was 72 hours.

### RNA short read sequencing

Poly(A)+ sequencing libraries were generated from total RNA using the NEBNext Ultra II Directional RNA Library Prep Kit (New England Biolabs) according to the manufacturer’s instruction, and sequenced on a NextSeq 500 device with single-end 76 cycles.

### De novo genome assembly

Raw unprocessed reads were assembled using *Canu* assembler (Koren et al. 2017) with an estimated genome size of 2.1gb and default parameters. This initial assembly was improved in a multi-step procedure. As a first step we mapped raw long reads using *minimap2* (Li 2018) and applied the *Racon* (Vaser et al. 2017) assembly polishing tool. In the next step, the *Racon*-polished genome was further improved with *Medaka* (https://nanoporetech.github.io/medaka) using again the raw long reads mapped with the *mini_align* script provided by the *Medaka* package, which also uses *minimap2*. For performance reasons, the polishing followed a recommendation in *Medaka’s* documentation and we first split the contigs into ten almost equally sized sets. Each set was processed using the subprogram *medaka consensus* and finally merged with *medaka stitch.* As a last polishing step, the result of *Medaka* was polished using the genomic short reads with *Polca* (Zimin & Salzberg 2020). Short reads were first trimmed and quality filtered using *bbduk (https://sourceforge.net/projects/bbmap/)* and filtered reads were used as input for *Polca*, which is based on *bwa-mem* read mapping and subsequent variant calling with *freebayes* (Garrison & Marth 2012). For evaluation of assembly quality and polishing effects, we applied the quality assessment tools *QUAST* (Mikheenko et al. 2018) and *BUSCO* (Simão et al. 2015) for estimation of genome completeness. At the final stage of the assembly process we performed contamination screening of the polished assembly based on *Kraken2* (Wood et al. 2019) with database option *standard* augmented with genomes *Mesocricetus auratus* and *Mus musculus,* retaining those contigs classified either as one of these two species or unclassified. Contigs classified as bacteria were removed and those classified as human were further analyzed using *BLASTN* (Altschul et al. 1990) with database *nt* to eliminate possible contamination with human genetic material. For every contig we summed up the bitscores per taxid for all hits with e-values below 1e-25 and assigned the species with the highest summed score. All contigs with hits to order *Rodentia* and or without any hits passing the threshold remained in the final assembly. The versions and options for all tools in the bioinformatics toolchain are given in Supplemental Table S4.

### Genome annotation

RNA-Seq reads were quality trimmed and adapter sequences were removed with *Cutadapt* (Martin 2011) and filtered reads were mapped to the final polished assembly using the mapper *STAR* (Dobin et al. 2013). The mapped reads, together with closely related reference genomes and annotations of *Mus musculus* (GRCm38.102), *Rattus norvegicus* (Rnor_6.0.102) and *Mesocricetus auratus* (MesAur1.0.100) - obtained from ENSEMBL - were used a input for the hybrid genome annotation tool *GeMoMa* (Keilwagen et al. 2016, 2018) to predict gene loci. The mapped RNA-Seq reads were also used in a subsequent prediction of 5’ and 3’ UTRs. Finally, resulting gff files were converted to gtf format using *GffRead* (Pertea & Pertea 2020) and augmented with the original gene name(s) of the associated gene from the reference genomes with a custom Python script. Afterwards the annotation was cleaned according to the following scheme: If transcripts annotated for a single locus matched different gene names, only transcripts associated to the same gene name as the highest scoring (*GeMoMa* score) transcript for this locus were retained. In a second step, if the same gene name was associated with multiple annotated loci, only the locus with the higher top score was retained. In another post-processing step we removed exons shared by multiple genes as these fusions were artifacts introduced by GeMoMa’s UTR inference. Finally, for transcripts with identical exon boundaries, all but the one with the longest CDS were removed with the script *agat_sp_fix_features_locations_duplicated.pl* from the AGAT toolkit (Dainat et al. 2021).

Finally, as we observed that annotated 3’ UTRs were frequently too short, we extended them by a constant number of 1000 bp whenever their distance to the next annotated feature (same or opposite strand) was at least 3000 bp.

## Acknowledgments

Supported by the Project “Virological and immunological determinants of COVID-19 pathogenesis – lessons to get prepared for future pandemics (KA1-Co-02 “COVIPA”)”, a grant from the Helmholtz Association’s Initiative and Networking Fund.

The authors thank Elisabeth Kirst, Jeannine Wilde and Madlen Sohn for sequencing support.

## Data Availability

The genomic sequencing data underlying this article are available in the European Nucleotide Archive (ENA) and can be accessed with accession numbers ERR6740384, ERR6740385 (Illumina) and ERR6797440 (ONT). The accession numbers for the RNA-Seq raw reads are ERR6752847 (pr-d0-lung-1), ERR6752848 (pr-d2-lung-1), ERR6752849 (pr-d2-lung-2) and ERR6752850 (pr-d3-lung-2).

The assembled genome together with annotation has been uploaded to figshare (https://doi.org/10.6084/m9.figshare.16695457 - ENA submission pending).

## Supplementary Material

**Table S1:** QUAST results for assembly and after different polishing steps

| Assembly | Canu | Canu +<br>Racon | Canu +<br>Racon +<br>Medaka | Canu +<br>Racon +<br>Medaka +<br>Polca | Canu +<br>Racon +<br>Medaka +<br>Polca +<br>Decont |
| --- | --- | --- | --- | --- | --- |
| # contigs ($\geq 0$ bp) | 2,112 | 2,109 | 2,108 | 2,108 | 2,078 |
| # contigs ($\geq 1000$ bp) | 2,112 | 2,109 | 2,108 | 2,108 | 2,078 |
| # contigs ($\geq 5000$ bp) | 2,109 | 2,108 | 2,108 | 2,108 | 2,078 |
| # contigs ($\geq 10000$ bp) | 2,108 | 2,107 | 2,107 | 2,107 | 2,077 |
| # contigs ($\geq 25000$ bp) | 2,106 | 2,105 | 2,105 | 2,105 | 2,075 |
| # contigs ($\geq 50000$ bp) | 2,086 | 2,085 | 2,085 | 2,085 | 2,055 |
| Total length ($\geq 0$ bp) | 2,381,232,474 | 2,386,917,682 | 2,388,794,081 | 2,388,036,258 | 2,384,223,288 |
| Total length ($\geq 1000$ bp) | 2,381,232,474 | 2,386,917,682 | 2,388,794,081 | 2,388,036,258 | 2,384,223,288 |
| Total length ($\geq 5000$ bp) | 2,381,220,014 | 2,386,913,209 | 2,388,794,081 | 2,388,036,258 | 2,384,223,288 |
| Total length ($\geq 10000$ bp) | 2,381,210,169 | 2,386,903,358 | 2,388,788,685 | 2,388,030,857 | 2,384,217,887 |
| Total length ($\geq 25000$ bp) | 2,381,168,154 | 2,386,861,164 | 2,388,746,456 | 2,387,988,633 | 2,384,175,663 |
| Total length ($\geq 50000$ bp) | 2,380,358,702 | 2,386,050,351 | 2,387,934,087 | 2,387,176,465 | 2,383,363,495 |
| # contigs | 2,112 | 2,109 | 2,108 | 2,108 | 2,078 |
| Largest contig | 79,389,475 | 79,566,412 | 79,616,872 | 79,594,593 | 79,594,593 |
| Total length | 2,381,232,474 | 2,386,917,682 | 2,388,794,081 | 2,388,036,258 | 2,384,223,288 |
| GC (%) | 41.31 | 41.27 | 41.33 | 41.32 | 41 |
| N50 | 25,694,176 | 25,762,496 | 25,788,438 | 25,775,705 | 25,775,705 |
| N75 | 7,300,843 | 7,317,947 | 7,325,048 | 7,322,524 | 7,322,524 |
| L50 | 30 | 30 | 30 | 30 | 30 |
| L75 | 73 | 73 | 73 | 73 | 73 |
| Illumina reads |  |  |  |  |  |
| # total reads | 1,354,820,496 | 1,354,379,523 | 1,354,069,760 | 1,354,067,462 | 1,354,038,936 |
| # left | 676,472,544 | 676,472,544 | 676,472,544 | 676,472,544 | 676,472,544 |
| # right | 676,472,544 | 676,472,544 | 676,472,544 | 676,472,544 | 676,472,544 |
| Mapped (%) | 99.73 | 99.75 | 99.75 | 99.75 | 99.75 |
| Properly paired (%) | 98.75 | 98.94 | 98.99 | 98.93 | 99.02 |
| Avg. coverage depth | 80 | 80 | 80 | 80 | 80 |
| Coverage $\geq 1x$ (%) | 99.66 | 99.57 | 98.94 | 98.96 | 98.99 |
| # N's per 100 kbp | 0 | 0 | 0 | 0 | 0 |
| ONT reads |  |  |  |  |  |
| # total reads | 4,265,943 | 4,206,546 | 4,154,967 | 4,153,904 | 4,133,827 |
| Mapped (%) | 99.73 | 99.73 | 99.74 | 99.74 | 99.74 |
| Properly paired (%) | 0 | 0 | 0 | 0 | 0 |
| Avg. coverage depth | 34 | 34 | 34 | 34 | 34 |
| Coverage >= 1x (%) | 99.99 | 99.99 | 99.98 | 99.98 | 99.99 |
| # N's per 100 kbp | 0.00 | 0.00 | 0.00 | 0.00 | 0 |

**Table S2:** BUSCO results for assembly after different polishing steps. Decontamination of POLCA cleaned assembly did not affect BUSCO statistics.

| **Assembly** | **Canu** | **Canu + Racon** | **Canu + Racon + Medaka** | **Canu + Racon + Medaka + POLCA (+ Decont)** |
| --- | --- | --- | --- | --- |
| Complete BUSCOs (C) | 10944 | 11857 | 12248 | 12692 |
| Complete and single-copy BUSCOs (S) | 10709 | 11616 | 12041 | 12467 |
| Complete and duplicated BUSCOs (D) | 235 | 241 | 207 | 225 |
| Fragmented BUSCOs (F) | 626 | 440 | 370 | 267 |
| Missing BUSCOs (M) | 2228 | 1501 | 1180 | 839 |
| Total BUSCO groups searched | 13798 | 13798 | 13798 | 13798 |

**Table S3:** Mapping statistics of RNA-Seq reads

| **Sample** | **pr-d0-lung-1** | **pr-d2-lung-1** | **pr-d2-lung-2** | **pr-d3-lung-2** |
| --- | --- | --- | --- | --- |
| Number of reads | 24,523,437 | 22,663,402 | 13,547,346 | 31,230,371 |
| Number of input reads (after quality filtering) | 23,667,474 | 22,230,068 | 13,222,122 | 30,107,224 |
| Average input read length | 74 | 74 | 74 | 74 |
| **UNIQUE READS:** |  |  |  |  |
| Uniquely mapped reads number | 18,414,197 | 18,347,689 | 11,543,772 | 21,555,625 |
| Uniquely mapped reads % | 77.80% | 82.54% | 87.31% | 71.60% |
| Average mapped length | 73.82 | 74.06 | 74.03 | 73.96 |
| Number of splices: Total | 2,714,011 | 3,019,367 | 1,750,395 | 3,522,768 |
| Number of splices: Annotated (sjdb) | 0 | 0 | 0 | 0 |
| Number of splices: GT/AG | 2,700,976 | 3,005,810 | 1,741,136 | 3,508,197 |
| Number of splices: GC/AG | 10,524 | 10,827 | 7,509 | 11,828 |
| Number of splices: AT/AC | 1,332 | 1,521 | 949 | 1,387 |
| Number of splices: Non-canonical | 1,179 | 1,209 | 801 | 1,356 |
| Mismatch rate per base, % | 0.17% | 0.15% | 0.15% | 0.16% |
| Deletion rate per base | 0.01% | 0.01% | 0.01% | 0.01% |
| Deletion average length | 1.51 | 1.48 | 1.51 | 1.48 |
| Insertion rate per base | 0.01% | 0.00% | 0.01% | 0.00% |
| Insertion average length | 1.39 | 1.35 | 1.36 | 1.35 |
| MULTI-MAPPING READS: |  |  |  |  |
| Number of reads mapped to multiple loci | 3,939,690 | 3,293,844 | 1,200,247 | 5,523,145 |
| % of reads mapped to multiple loci | 16.65% | 14.82% | 9.08% | 18.34% |
| Number of reads mapped to too many loci | 553,871 | 295,740 | 200,857 | 1,985,491 |
| % of reads mapped to too many loci | 2.34% | 1.33% | 1.52% | 6.59% |
| UNMAPPED READS: |  |  |  |  |
| Number of reads unmapped: too many mismatches | 0 | 0 | 0 | 0 |
| % of reads unmapped: too many mismatches | 0.00% | 0.00% | 0.00% | 0.00% |
| Number of reads unmapped: too short | 540,151 | 182,355 | 199,639 | 422,951 |
| % of reads unmapped: too short | 2.28% | 0.82% | 1.51% | 1.40% |
| Number of reads unmapped: other | 219,565 | 110,440 | 77,607 | 620,012 |
| % of reads unmapped: other | 0.93% | 0.50% | 0.59% | 2.06% |
| CHIMERIC READS: |  |  |  |  |
| Number of chimeric reads | 0 | 0 | 0 | 0 |
| % of chimeric reads | 0.00% | 0.00% | 0.00% | 0.00% |

**Table S4:**
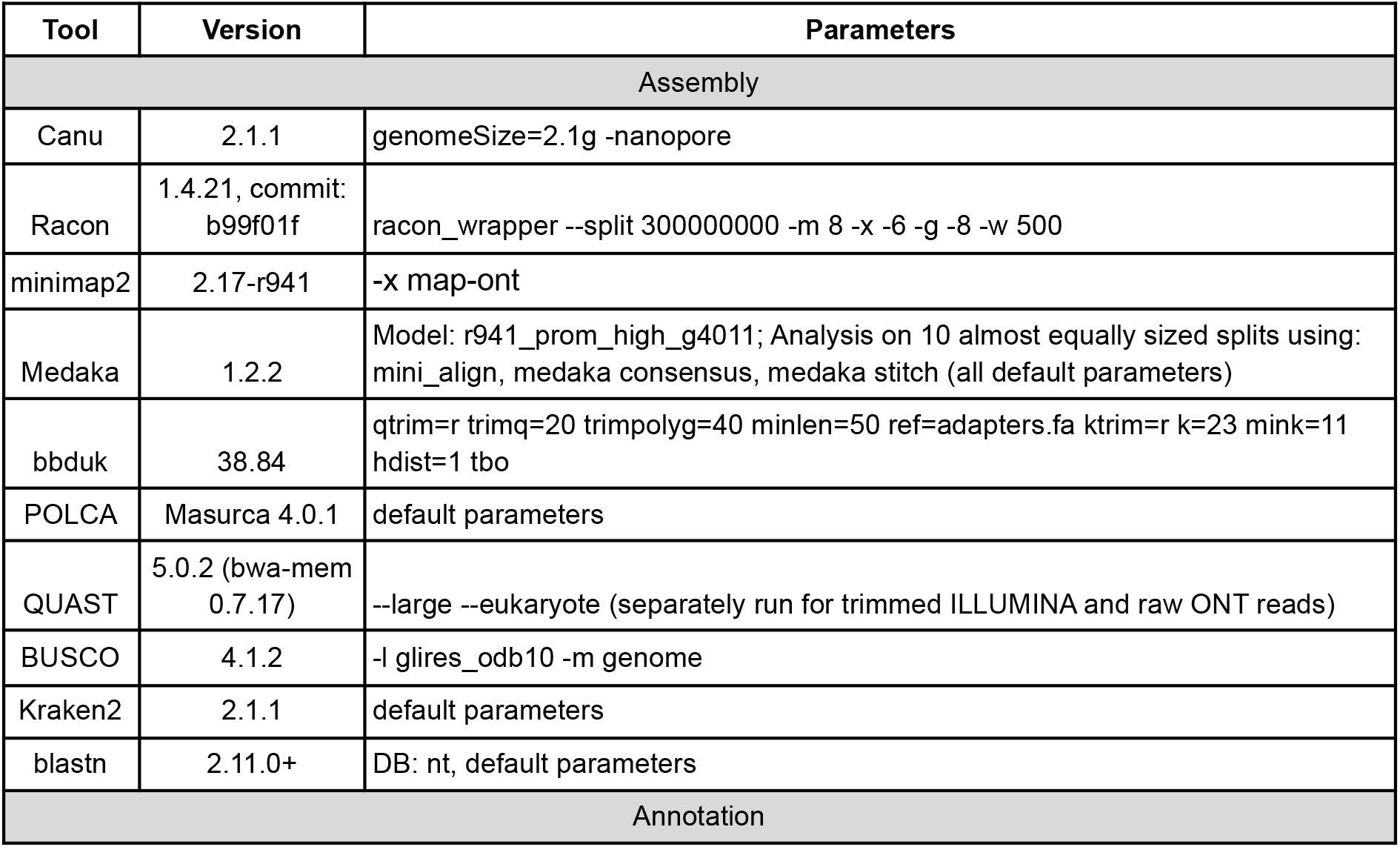
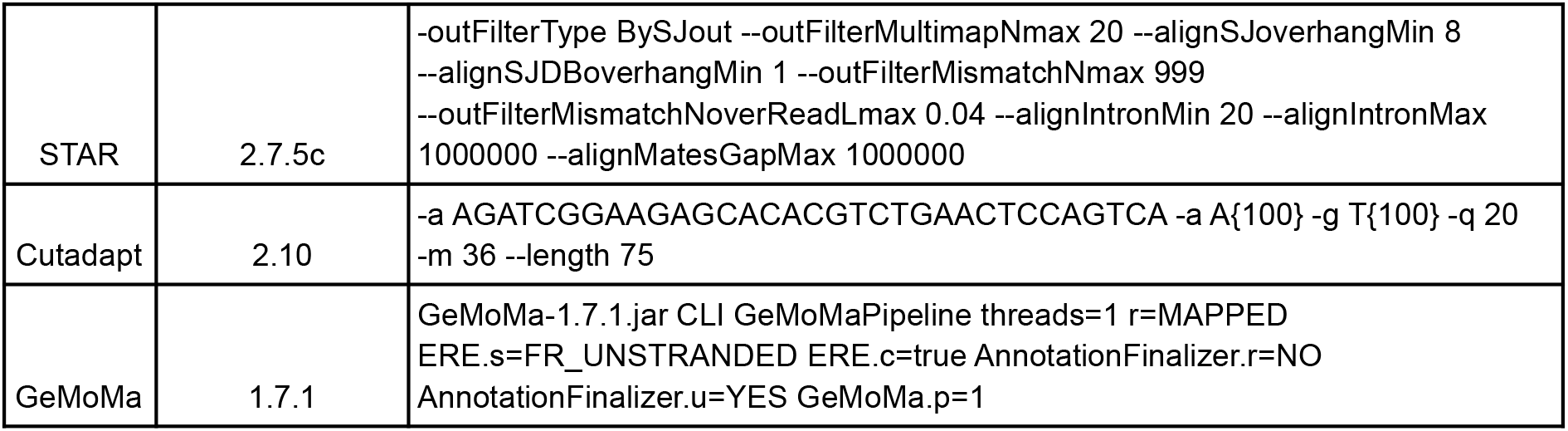
Overview tools, versions and commands:

## References

Altschul SF, Gish W, Miller W, Myers EW, Lipman DJ. 1990. Basic local alignment search tool. J. Mol. Biol. 215:403–410. doi: 10.1016/S0022-2836(05)80360-2.

Bao L et al. 2020. The pathogenicity of SARS-CoV-2 in hACE2 transgenic mice. Nature. 583:830–833. doi: 10.1038/s41586-020-2312-y.

Bertzbach LD et al. 2021. SARS-CoV-2 infection of Chinese hamsters (*Cricetulus griseus*) reproduces COVID-19 pneumonia in a well-established small animal model. Transbound. Emerg. Dis. 68:1075–1079. doi: 10.1111/tbed.13837.

Chan JF-W et al. 2020. Simulation of the Clinical and Pathological Manifestations of Coronavirus Disease 2019 (COVID-19) in a Golden Syrian Hamster Model: Implications for Disease Pathogenesis and Transmissibility. Clin. Infect. Dis. ciaa325. doi: 10.1093/cid/ciaa325.

Dainat J, Hereñú D, Pascal-Git. 2021. NBISweden/AGAT: AGAT-v0.8.0. Zenodo doi: 10.5281/ZENODO.3552717.

Deng W et al. 2020. Ocular conjunctival inoculation of SARS-CoV-2 can cause mild COVID-19 in rhesus macaques. Nat. Commun. 11:4400. doi: 10.1038/s41467-020-18149-6.

Devaux CA, Pinault L, Osman IO, Raoult D. 2021. Can ACE2 Receptor Polymorphism Predict Species Susceptibility to SARS-CoV-2? Front. Public Health. 8:608765. doi: 10.3389/fpubh.2020.608765.

Dinnon KH et al. 2020. A mouse-adapted model of SARS-CoV-2 to test COVID-19 countermeasures. Nature. 586:560–566. doi: 10.1038/s41586-020-2708-8.

Dobin A et al. 2013. STAR: ultrafast universal RNA-seq aligner. Bioinformatics. 29:15–21. doi: 10.1093/bioinformatics/bts635.

Gantier MP. 2021. Animal models of COVID-19 hyper-inflammation. Respirol. Carlton Vic. 26:222–224. doi: 10.1111/resp.13997.

Garrison E, Marth G. 2012. Haplotype-based variant detection from short-read sequencing. ArXiv12073907 Q-Bio. http://arxiv.org/abs/1207.3907 (Accessed April 28, 2021).

Gu H et al. 2020. Adaptation of SARS-CoV-2 in BALB/c mice for testing vaccine efficacy. Science. 369:1603–1607. doi: 10.1126/science.abc4730.

Hassan AO et al. 2020. A SARS-CoV-2 Infection Model in Mice Demonstrates Protection by Neutralizing Antibodies. Cell. 182:744–753.e4. doi: 10.1016/j.cell.2020.06.011.

Imai M et al. 2020. Syrian hamsters as a small animal model for SARS-CoV-2 infection and countermeasure development. Proc. Natl. Acad. Sci. 202009799. doi: 10.1073/pnas.2009799117.

Keilwagen J et al. 2016. Using intron position conservation for homology-based gene prediction. Nucleic Acids Res. 44:e89–e89. doi: 10.1093/nar/gkw092.

Keilwagen J, Hartung F, Paulini M, Twardziok SO, Grau J. 2018. Combining RNA-seq data and homology-based gene prediction for plants, animals and fungi. BMC Bioinformatics. 19:189. doi: 10.1186/s12859-018-2203-5.

Kim Y-I et al. 2020. Infection and Rapid Transmission of SARS-CoV-2 in Ferrets. Cell Host Microbe. 27:704–709.e2. doi: 10.1016/j.chom.2020.03.023.

Koren S et al. 2017. Canu: scalable and accurate long-read assembly via adaptive *k*-mer weighting and repeat separation. Genome Res. 27:722–736. doi: 10.1101/gr.215087.116.

Kreye J et al. 2020. A Therapeutic Non-self-reactive SARS-CoV-2 Antibody Protects from Lung Pathology in a COVID-19 Hamster Model. Cell. 183:1058–1069.e19. doi: 10.1016/j.cell.2020.09.049.

Lee C-Y, Lowen AC. 2021. Animal models for SARS-CoV-2. Curr. Opin. Virol. 48:73–81. doi: 10.1016/j.coviro.2021.03.009.

Li H. 2018. Minimap2: pairwise alignment for nucleotide sequences Birol, I, editor. Bioinformatics. 34:3094–3100. doi: 10.1093/bioinformatics/bty191.

Lu S et al. 2020. Comparison of nonhuman primates identified the suitable model for COVID-19. Signal Transduct. Target. Ther. 5:157. doi: 10.1038/s41392-020-00269-6.

Martin M. 2011. Cutadapt removes adapter sequences from high-throughput sequencing reads. EMBnet.journal. 17:10. doi: 10.14806/ej.17.1.200.

Mikheenko A, Prjibelski A, Saveliev V, Antipov D, Gurevich A. 2018. Versatile genome assembly evaluation with QUAST-LG. Bioinformatics. 34:i142–i150. doi: 10.1093/bioinformatics/bty266.

Muñoz-Fontela C et al. 2020. Animal models for COVID-19. Nature. 586:509–515. doi: 10.1038/s41586-020-2787-6.

Osterrieder N et al. 2020. Age-Dependent Progression of SARS-CoV-2 Infection in Syrian Hamsters. Viruses. 12:E779. doi: 10.3390/v12070779.

Pach S et al. 2020. ACE2-Variants Indicate Potential SARS-CoV-2-Susceptibility in Animals: An Extensive Molecular Dynamics Study. Pharmacology and Toxicology doi: 10.1101/2020.05.14.092767.

Pertea G, Pertea M. 2020. GFF Utilities: GffRead and GffCompare. F1000Research. 9:304. doi: 10.12688/f1000research.23297.1.

Richard M et al. 2020. SARS-CoV-2 is transmitted via contact and via the air between ferrets. Nat. Commun. 11:3496. doi: 10.1038/s41467-020-17367-2.

Shi J et al. 2020. Susceptibility of ferrets, cats, dogs, and other domesticated animals to SARS–coronavirus 2. Science. 368:1016–1020. doi: 10.1126/science.abb7015.

Sia SF et al. 2020. Pathogenesis and transmission of SARS-CoV-2 in golden hamsters. Nature. 583:834–838. doi: 10.1038/s41586-020-2342-5.

Simão FA, Waterhouse RM, Ioannidis P, Kriventseva EV, Zdobnov EM. 2015. BUSCO: assessing genome assembly and annotation completeness with single-copy orthologs. Bioinformatics. 31:3210–3212. doi: 10.1093/bioinformatics/btv351.

Sun J et al. 2020. Generation of a Broadly Useful Model for COVID-19 Pathogenesis, Vaccination, and Treatment. Cell. 182:734–743.e5. doi: 10.1016/j.cell.2020.06.010.

Trimpert J et al. 2021. Development of safe and highly protective live-attenuated SARS-CoV-2 vaccine candidates by genome recoding. Cell Rep. 36:109493. doi: 10.1016/j.celrep.2021.109493.

Trimpert J et al. 2020. The Roborovski Dwarf Hamster Is A Highly Susceptible Model for a Rapid and Fatal Course of SARS-CoV-2 Infection. Cell Rep. 33:108488. doi: 10.1016/j.celrep.2020.108488.

Vaser R, Sović I, Nagarajan N, Šikić M. 2017. Fast and accurate de novo genome assembly from long uncorrected reads. Genome Res. 27:737–746. doi: 10.1101/gr.214270.116.

Veenhuis RT, Zeiss CJ. 2021. Animal Models of COVID-19 II. Comparative Immunology. ILAR J. ilab010. doi: 10.1093/ilar/ilab010.

Wilson DE, Mittermeier RA, Cavallini P, eds. 2009. Handbook of the mammals of the world - Volume 7: Rodents II. Lynx Edicions : Conservation International: IUCN: Barcelona.

Wood DE, Lu J, Langmead B. 2019. Improved metagenomic analysis with Kraken 2. Genome Biol. 20:257. doi: 10.1186/s13059-019-1891-0.

Wu C et al. 2020. In Silico Analysis of Intermediate Hosts and Susceptible Animals of SARS-CoV-2. doi: 10.26434/chemrxiv.12057996.v1.

Yu P et al. 2020. Age-related rhesus macaque models of COVID-19. Anim. Models Exp. Med. 3:93–97. doi: 10.1002/ame2.12108.

Zhai C et al. 2021. Roborovski hamster *(Phodopus roborovskii)* strain SH101 as a systemic infection model of SARS-CoV-2. Virulence. 12:2430–2442. doi: 10.1080/21505594.2021.1972201.

Zimin AV, Salzberg SL. 2020. The genome polishing tool POLCA makes fast and accurate corrections in genome assemblies Ouzounis, CA, editor. PLOS Comput. Biol. 16:e1007981. doi: 10.1371/journal.pcbi.1007981.

